# Two cytochrome P450 epoxidases mediate juvenile hormone biosynthesis in *Drosophila melanogaster*

**DOI:** 10.1101/2025.10.24.684410

**Authors:** Daiki Fujinaga, Yuya Ohhara, Naoki Okamoto, Hannah Chu, Kerry E. Mauck, Naoki Yamanaka

## Abstract

Juvenile hormones (JHs) mediate various biological processes such as development and reproduction in insects. Although pleiotropic functions of JHs are well investigated in the fruit fly *Drosophila melanogaster*, their biosynthetic mechanisms are less well understood, partly because many JH biosynthetic enzymes still remain unidentified in this important model species. Here we report that two cytochrome P450 (CYP) epoxidases mediate JH biosynthesis in *D. melanogaster*. In addition to previously reported Cyp6g2, a second epoxidase, Cyp6a13, also functions in the corpus allatum, the major JH biosynthetic endocrine gland. Combined mutations of the genes encoding these enzymes cause developmental and reproductive defects, which can be rescued by JH application. JH biosynthetic functions of these genes were further confirmed by using a heterologous expression system and *ex vivo* tissue culture. Collectively, our results indicate that these two CYP epoxidases function cooperatively to mediate JH biosynthesis in *D. melanogaster*.

**Highlights:**

- *Drosophila* Cyp6g2 and Cyp6a13 are highly expressed in the corpus allatum.
- Combined knockout mutants of the CYP genes exhibit developmental and reproductive defects.
- Mutant phenotypes were rescued by feeding juvenile hormones.
- Both Cyp6g2 and Cyp6a13 catalyze epoxidation of juvenile hormone precursors.

## 1. Introduction

Juvenile hormones (JHs) are aliphatic sesquiterpenoids that regulate diverse biological processes in insects, including development and reproduction (Jindra et al., 2013; Jindra and Noriega, 2026). JH acts as a “status quo” hormone that maintains the juvenile form across many insect orders during early postembryonic development (Riddiford, 2020), and a decline in the hemolymph JH titer triggers metamorphosis (Jindra, 2019). Ablation of the corpora allata (CA), the JH-producing glands, causes precocious metamorphosis, whereas ectopic application of JH analogs such as methoprene induces supernumerary larval or nymphal molts in various species (Konopova et al., 2011; Kayukawa et al., 2012). JH is thus considered as a key regulator that prevents metamorphosis (Jindra et al., 2013). Intriguingly, JH is also produced at the onset of metamorphosis in holometabolous insects (Jindra, 2019). Disruption of JH receptors causes pupation defects (Konopova and Jindra, 2007; Abdou et al., 2011), suggesting that JH contributes to the proper initiation of metamorphosis, potentially by preventing premature adult differentiation (Martín et al., 2026).

The biosynthetic pathway of JH is well characterized and divided into early and late steps (Bellés et al., 2005). In the early steps, known collectively as the mevalonate pathway, farnesyl pyrophosphate (FPP) is synthesized. In late steps, FPP is first converted into farnesoic acid (FA) by a phosphatase and reductases (Noriega, 2014). The sequence of the final JH biosynthetic steps differs among insect taxa. In Lepidoptera, FA is epoxidized to JH acid (JHA) by a cytochrome P450 (CYP) epoxidase and then methylated by JHA methyltransferase (Jhamt) to produce JH (Daimon et al., 2012; Shinoda et al., 2003). In contrast, in most other insect orders, FA is first methylated by Jhamt to form methyl farnesoate (MF), which is subsequently epoxidized to yield JH (Helvig et al., 2004; Defelipe et al., 2011). Once synthesized, JH regulates gene expression in peripheral tissues by interacting with its intracellular receptor Methoprene-tolerant (Met). In the fruit fly *Drosophila melanogaster*, this function is shared by its paralog, germ-cell expressed (gce) (Konopova et al., 2007; Jindra et al., 2015).

The molecular mechanisms of JH-mediated developmental regulation have been extensively investigated in *D. melanogaster* using genetic tools such as the Gal4-UAS system. Although genetically allatectomized *Drosophila* larvae (also referred to as CA-ablated or CAX larvae) do not undergo precocious metamorphosis, they exhibit premature differentiation of some tissues, such as the eye disks and fat body (Liu et al., 2009; Riddiford et al., 2010; Zhang et al., 2022). Both CAX and JH receptor knockout larvae remain viable until pupariation, but they typically die around the time of head eversion, failing to complete pupation (Riddiford et al., 2010; Abdou et al., 2011). In adults, JH functions as a pleiotropic hormone. It acts as a gonadotropin by promoting vitellogenesis in females and accessory gland protein synthesis in males (Wilson et al., 2003; Santos et al., 2019). Recent studies further implicate JH in the regulation of courtship behavior, sleep, and aging (Wijesekera et al., 2016; Wu et al., 2018; Yamamoto et al., 2013).

In contrast to the well-characterized downstream effects of JH signaling, the regulation of JH synthesis remains relatively understudied in *D. melanogaster*. Notably, several enzymes catalyzing reactions in the late steps have yet to be fully identified (Fig. 1). For example, mutants lacking *jhamt* do not exhibit the pupal lethal phenotype observed in *Met* and *gce* double mutants (Abdou et al., 2011; Wen et al., 2015), suggesting the existence of other methyltransferase(s) that compensate for the loss of Jhamt. Moreover, a canonical CYP epoxidase is not conserved in higher dipterans. A recent study identified Cyp6g2 as an epoxidase in *D. melanogaster* (Jia et al., 2024). This enzyme synthesizes not only JH III, the predominant form of JH in insects, but also JH III bisepoxide (JHB3), a higher dipteran-specific JH (Richard et al., 1989). However, the existence of additional epoxidases remains plausible, as *Cyp6g2* disruption does not cause severe developmental defects as compared to JH receptor knockouts (Abdou et al. 2011; Jia et al. 2024).

**Figure 1.**
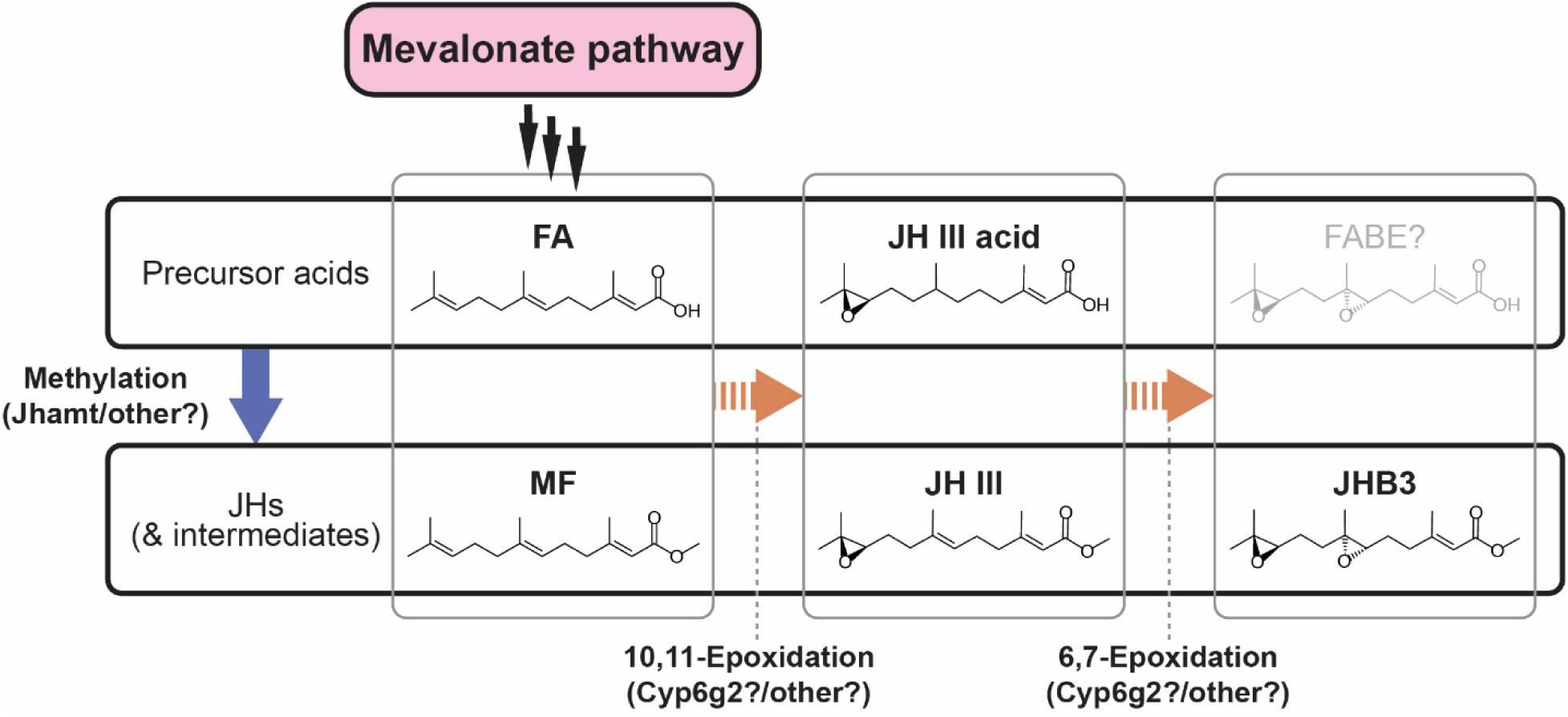
Last steps of the JH biosynthetic pathway in *Drosophila melanogaster*. Farnesoic acid (FA) is converted into juvenile hormone III (JH III) and JH-bisepoxide (JHB3) in the corpus allatum. Cyp6g2 predominantly epoxidizes FA into JH III acid, rather than methyl farnesoate (MF) into JH III (Jia et al., 2024). Jhamt converts both FA and JH III acid into MF and JH III, respectively (Niwa et al., 2008). Although FA bisepoxide (FABE) has been hypothesized as an intermediate of JHB3 synthesis (Moshitzky and Applebaum, 1995), it has not been detected in *D. melanogaster*.

In the present study, we investigated CYP genes that are highly expressed in the CA in *D. melanogaster* and identified a second epoxidase, Cyp6a13. Combined knockouts of *jhamt*, *Cyp6g2*, and *Cyp6a13* caused severe developmental defects similar to those observed in *Met* and *gce* double mutants. These mutants also showed reduced egg production, a phenotype commonly associated with JH deficiency. Furthermore, we confirmed the enzymatic functions of these genes using a heterologous expression system and *ex vivo* tissue culture. Collectively, our results indicate that Cyp6g2 and Cyp6a13 function cooperatively to mediate JH biosynthesis in *D. melanogaster*.

## 2. Materials and methods

### 2.1. Flies

Flies were raised at 25°C under a 12 h-light and 12 h-dark photocycle on a standard cornmeal diet (Fujinaga et al., 2025). The control strain *w^1118^* and transgenic flies were obtained from the Bloomington *Drosophila* Stock Center (BDSC), Vienna *Drosophila* Resource Center (VDRC), Tzumin Lee, Michael E. Adams, and Michael B. O’Connor, as listed in Table S1. Complementary DNA (cDNA) fragments of *jhamt*, *Cyp6g2*, and *Cyp6a13* were obtained from cDNA clones AT13581, IP03155, and LD25139, respectively, from the *Drosophila* Genomic Resource Center (DGRC) and cloned into the *pUAST* vector (Table S2) to generate UAS constructs. Transgenic flies were generated by BestGene Inc. Null mutant flies were generated by deleting most or all of the coding sequences of the target genes using the CRISPR-Cas9 system. Mutagenesis of *Met*, *gce*, and *Cyp6a13* was performed following the protocol developed by Kondo and Ueda (2013), with slight modifications as described in our previous study (Fujinaga et al., 2025). Briefly, two guide RNA (gRNA) target sequences for each genomic locus were inserted into *pBFv-U6.2* vectors using oligonucleotide pairs listed in Table S3, and a plasmid with two gRNA sequences was constructed for each gene. For the *Met* and *gce* mutant strains, embryonic injection of the plasmids and generation of G1 mutants were performed by BestGene Inc. For the *Cyp6a13* mutant strain, a transgenic strain carrying the plasmid at the *attP2* site was generated by BestGene Inc. The obtained *U6-gRNA* flies were crossed with *nos-Cas9* flies to induce mutagenesis in germ cells. Mutations were confirmed using primers designed outside the target sites (Table S3). Gene disruption mutants of *jhamt* and *Cyp6g2* were generated by WellGenetics Inc., using CRISPR-based homologous recombination to insert fluorescent protein-coding sequences. All mutants were backcrossed with the *w^1118^* control strain for at least four generations to minimize off-target effects and standardize genetic backgrounds.

### 2.2. Cell lines

Human embryonic kidney (HEK) 293 cells were obtained from Michael E. Adams. Cells were maintained in 75 cm^2^ flasks (VWR) in a humidified incubator at 37°C and 5% CO_2_ in Dulbecco’s Modified Eagle’s Medium (DMEM) with 4.5 mg/mL glucose, L-glutamine, and sodium pyruvate (Corning, #10-013-CV) containing 10% fetal bovine serum (Corning, #10082-139) and 1% penicillin-streptomycin (Thermo Fisher Scientific, #15-140-122).

### 2.3. Transcriptome analysis using public RNA sequencing data

Public RNA sequencing data (GSE229077) were obtained from the Gene Expression Omnibus (Roger et al., 2023). Reads were aligned to the *Drosophila* genome (BDGP6.46, Ensembl) using Hisat2 (Kim et al., 2019). Aligned reads per gene were counted using SAMtools (Danecek et al., 2021) and featureCount (Liao et al., 2014). Reads per kilobase per million (RPKM) were calculated based on total reads and are listed in Table S4. Genes with expression levels above 20 RPKM were considered to be highly expressed.

### 2.4. Visualization of *jhamt-LexA* expression patterns

Wandering larvae and three-day post-eclosion adult males expressing *jhamt-LexA*-driven *LexAop-mCD8::GFP* were chilled on ice for 10 min. GFP expression patterns in the whole body were observed using a SteREO Discovery.V12 microscope (Zeiss). Detailed expression patterns of *jhamt-LexA* were visualized by immunostaining. Tissues were dissected in phosphate-buffered saline (PBS) (Fisher Scientific, #BP399-1), fixed with 4% paraformaldehyde (PFA) (Electron Microscopy Sciences, #157-8) in PBS containing 0.1% TritonX-100 (4% PFA/PBSTx) for 20 minutes at room temperature (RT), and washed multiple times with PBSTx. Tissues were blocked with 5% normal goat serum (NGS) (Sigma-Aldrich, G9023) in PBSTx for at least one hour at RT, incubated overnight at 4°C with chicken anti-GFP antibody (1:500, Abcam, #ab13970) in PBSTx with 5% NGS, and then washed multiple times with PBSTx. They were then incubated for two hours at RT with goat anti-chicken IgG conjugated to Alexa Fluor 488 (Life Technologies, #A-11039) in PBSTx with 5% NGS, followed by additional PBSTx washes. DNA was stained with Hoechst 33342 (1:2,000, Life Technologies, #H3570) for 30 min at RT. After washing, tissues were mounted in Vectashield H-1000 (Vector Laboratories, #H-1000-10) and observed using a Zeiss Axio Imager M2 equipped with ApoTome.2 (Zeiss).

### 2.5. Total RNA extraction and quantitative reverse transcription PCR

Total RNA was extracted from tissues using TRIzol reagent (Thermo Fisher Scientific, #15596018), and purified with the RNeasy Mini Kit (Qiagen, #74106) following the manufacturers’ instructions. cDNA was synthesized from purified RNA using PrimeScript RT Master Mix (Takara Bio, #RR036A). Quantitative reverse transcription (qRT)-PCR was performed on a CFX Connect Real-Time PCR Detection System (Bio-Rad) using AzuraView™ GreenFast qPCR Blue Mix LR (Azura Genomics, AZ-2320). Serial dilutions of plasmids containing target sequences were used as standards for absolute quantification. Transcript levels of mRNA were normalized to the internal control gene, *rp49*, as in Okamoto et al., 2018. Primers are listed in Table S3.

### 2.6. RNA probe synthesis

Coding sequences of the target genes were PCR-amplified using T7- and SP6-tagged primers listed in Table S3. Digoxigenin (DIG)-labelled RNA was synthesized using the DIG RNA Labeling Kit (SP6/T7, Roche, #11175025910) following the manufacturer’s protocol. Following DNase treatment at 37°C for 15 min, labelled RNA was purified using the RNeasy Mini Kit. RNA was hydrolyzed at 60°C for 45 min in alkaline solution (80 mM sodium bicarbonate, 120 mM sodium carbonate, and 10 mM DTT), and the reaction was terminated with neutralization buffer (300 mM sodium acetate, 1% acetic acid, and 10 mM DTT). RNA probes were purified by ethanol precipitation and stored at -20°C.

### 2.7. Whole mount *in situ* hybridization

Wandering larvae of *w^1118^* were cut in half in PBS, and posterior segments were flipped to expose internal organs. Tissues were fixed in 4% PFA in PBS for 20 min, then washed three times with PBS containing 0.2% Tween 20 (PBST). Tissues were then rinsed three times with methanol. After washing with 90% methanol containing 50 mM EGTA, half the volume was replaced to 4% PFA three times, followed by fixation in 4% PFA in PBS for 20 min. Tissues were treated with 50 µg/mL Proteinase K (Roche, #3115828001) in PBST for 5 min, followed by fixation in 4% PFA in PBS for 20 min. After washing with PBST, tissues were incubated in the pre-hybridization buffer (50% formamide, saline sodium citrate, 100 µg/mL heparin, 100 µg/mL yeast RNA, 10 mM DTT, and 0.1% Tween 20) at 60°C for one hour. RNA probes (200 ng/mL for *jhamt* and *Cyp6g2*, 1,000 ng/mL for *Cyp6a13*) were denatured in the hybridization solution (10% dextran sulfate-containing pre-hybridization solution) at 80°C for 15 min, followed by chilling on ice for 5 min. The RNA probe solution was then applied to tissues for overnight incubation at 60°C. Tissues were washed six times for 30 min with the washing solution (50% formamide, saline sodium citrate, and 0.1% Tween 20), then blocked with the blocking solution (2% bovine serum albumin in PBST) for 30 min. Alkaline phosphatase-conjugated anti-DIG antibody (Roche, #11093274910) diluted in the blocking solution (1:3,000) was applied overnight at 4°C. Following three 20-min PBST washes, tissues were incubated in the alkaline phosphatase buffer (100 mM Tris-hydrochloride, 100 mM sodium chloride, 50 mM magnesium chloride, and 0.1% Tween 20, pH 9.5), followed by colorimetric development with nitro blue tetrazolium (Roche, #11383213001) and 5-bromo-4-chloro-3-indolyl phosphate (Roche, #11383221001) for one hour. After washing, tissues were permeated with glycerol, and the central nervous system-ring gland (RG) complexes were dissected and imaged using a SteREO Discovery.V12 microscope (Zeiss) and an Olympus BX53 microscope with a DP26 CCD camera.

### 2.8. Scoring of lethal stages

Eggs of each genotype were collected on a grape juice plate with yeast powder. Larvae were collected within two hours after hatching and reared on the standard cornmeal diet until 48 hours after hatching. Homozygous mutants were selected based on the absence of the balancer chromosome carrying GFP or RFP, followed by separation of males and females. Larvae were then transferred to new vials (8-25 larvae/vial) containing the standard cornmeal diet with or without specific concentrations of FA (Echeron Biosciences, #S-0151), MF (Echeron Biosciences, #S-0153), JH III (Echeron Biosciences, #S-0155), or JHB3 (Echeron Biosciences, #S-0157). Dead larvae and prepupae were counted six days after hatching, after which the size of the pupal cases was measured. Animals that underwent head eversion were scored as pupae. Dead pupae, early lethal adults, and normal adults were scored 10 days after hatching. The developmental lethal phenotype was scored as follows: larval lethal = 0, prepupal lethal = 1, pupal lethal = 2, early adult lethal = 3, and normal development = 4. The average score of each genotype was calculated as the developmental score, and it was plotted along with the *Kruppel homolog 1* (*Kr-h1*) expression level or the pupal case size for various genotypes using R. The correlation coefficient and *p*-value (Pearson’s test) for each plot were calculated using R.

### 2.9. Fecundity assay

Newly hatched larvae of selected genotypes were raised on the cornmeal diet until 48 hours after hatching. Larvae were then transferred to vials at a density of fewer than 40 larvae per 3 g of the diet and kept until eclosion. Five virgin females of each genotype were collected within six hours after eclosion and kept in the vials containing the cornmeal diet with or without 0.5 ppm MF or JH III. Twenty-four hours later, three *w^1118^* males were added for mating. After another 24 hours, flies were transferred to vials containing agarose-solidified grape juice with yeast powder. Flies were transferred into fresh vials every 24 hours, and the number of eggs in each vial was counted under a stereomicroscope.

### 2.10. Enzymatic conversion assay in HEK293 cells

Coding sequences of *jhamt*, *Cyp6g2*, and *Cyp6a13* for heterologous expression in HEK293 cells were obtained from cDNA clones AT13581, IP03155, and LD25139, respectively, as listed in Table S2, and cloned into the *pcDNA3.1(+)* vector. HEK293 cells at a density of 2 x 10^5^ cells in 2 mL of the culture medium were seeded on a 6-well clear flat bottom microplate (Bioland Scientific, #703003). After preincubation for 24 hours, 0.3 µg of the *jhamt*-encoding plasmid and 0.9 µg of CYP enzyme-encoding plasmids were transfected using Attractene transfection reagent (Qiagen, #301005) according to the manufacture’s instruction. The empty vector was transfected for control experiments. After 48 hours of incubation, 5 × 10^5^ cells were transferred into glass vials (Agilent, #5183-2068) containing S-adenosyl methionine (Millipore, #A7007; final concentration: 50 µM) and the esterase inhibitor Paraoxon (Fisher Scientific, #NC0811417; final concentration: 1 µM) in 720 µL of the FBS-free medium. Three hours after incubation, 7.2 µL of 1 mM FA was added, and the cells were further incubated for 18 hours. After mixing with 80 µL of acetonitrile, vials were stored at -20°C until extraction.

### 2.11. *Ex vivo* ring gland culture

Ten RGs of selected genotypes were dissected from wandering larvae and preincubated in 720 µL of Schneider’s *Drosophila* medium (Gibco, #21-720-024) in a glass vial for 30 min. After adding 7.2 µL of 1 mM FA to the culture medium, the RGs were incubated for four hours at 25°C. After incubation, 80 µL of acetonitrile was added, and the samples were stored at -20°C until extraction.

### 2.12. Hemolymph collection

Wandering third instar larvae were rinsed in PBS and dried on tissue paper. The cuticle was carefully torn to release the hemolymph onto a parafilm membrane. Ten µL of the hemolymph was collected and mixed with extraction solution (90 μl of 0.9% sodium chloride solution and 10 µL of acetonitrile) in a glass insert vial (Agilent, #5181-1270).

### 2.13. Sesquiterpenoid extraction

Extraction of sesquiterpenoids was performed based on the previous study (Sarro et al., 2021, adapted from Kai et al., 2018) with modifications. For hemolymph samples, 2 ng of citronellol (Millipore, #W230915) or 4 ng of deuterium-labelled JH III (JH III-d3) (LGC Ltd., #E589402) was added as an internal standard for gas chromatography-mass spectrometry (GC-MS) analysis or liquid chromatography-mass spectrometry (LC-MS) analysis, respectively. After 100 µL of hexane was added, the samples were vigorously mixed for 30 seconds. The samples were then allowed to sit for 5 min, followed by centrifugation at 1,000 x g for 5 min. The hexane layer was transferred to a new glass vial using a Hamilton syringe (Hamilton, #80600), and the extraction was repeated. For LC-MS analysis, the pooled hexane extract was concentrated by CentriVap concentrator (Labconco) to 100 µL. After adding 100 µL of methanol, the remaining hexane was evaporated. For extraction from the culture medium, 16 ng of citronellol or 4 ng of JH III-d3 was added. After adding 800 µL of hexane, the samples were vigorously mixed for 30 seconds. The hexane layer was then collected as described above.

### 2.14. Sesquiterpenoid determination

For developing the simultaneous extraction method, FA, MF, JH III, and JHB3 (100 ng each) were added to the extraction solution (400 or 720 µl of 0.9% sodium chloride solution and 400 or 80 µl of acetonitrile) in the glass vial. Compounds were extracted as described above, and the hexane extract was concentrated to 100 µl using CentriVap concentrator (Labconco). After adding 100 µl of methanol, the remaining hexane was evaporated. Extracted sesquiterpenoids were analyzed using a 1220 Infinity LC system (Agilent Technologies) with a Vydac 218TP C18 HPLC column (100 mm x 4.6 mm, 5 µm particle size, Grace). HPLC-grade acetonitrile and water were used as the liquid phase in a binary gradient flow (Pump A: 20% acetonitrile/80% water, Pump B: 100% acetonitrile). Ten µl of the extracted sample or authentic standards was injected, and absorbance at 217 nm was recorded for peak area analyses.

For biological samples, MF and JH III were detected and quantified using a Thermo Scientific Trace 1310 GC-MS equipped with an AI 1310 autosampler and a TSQ Duo triple quadrupole mass spectrometer (Thermo Scientific), following the method described previously (Sarro et al., 2021) with slight modifications. Detailed parameters for selected reaction monitoring were listed in Table S5. Authentic standards of MF and JH III were used to generate standard curves. Peak areas of the target compounds were normalized by the peak area of citronellol (10 ng/mL). JHB3 levels were determined using LC-MS at the UC Riverside Metabolomics Core based on the previous study (Ramirez et al., 2020) with modifications. Liquid chromatography separation was performed by an ultra-performance liquid chromatography system (ACQUITY UPLC I-Class PLUS, Waters) equipped with an ACQUITY UPLC BEH C18 column (2.1 × 100 mm, 1.7 µm, Waters). The column temperature was maintained at 35°C. Gradient separation was performed between 0.1% formic acid in water (mobile phase A) and 0.1% formic acid in acetonitrile (mobile phase B) at a flow rate of 0.3 mL/min. The gradient program was as follows: 4% B at 0 min; increase to 10% B in 2 min; increase to 80% B in 12 min and hold for 0.1 min; increase again to 100% B in 0.9 min and hold for another 0.1 min; then return to 4% B in 1.4 min; for a total run time of 16.5 min. Detection was performed using a Xevo TQ-XS mass spectrometry system (Waters) equipped with an electrospray ionization interface. The instrument was operated in multiple reaction monitoring mode as described in Table S6. Authentic standards of FA, JH III, and JHB3 were used to generate standard curves. Peak areas were normalized to the peak area of JH III-d3 (40 ng/mL).

### 2.15. Phylogenetic analysis

The unrooted maximum-likelihood phylogenetic tree was generated using MEGAX (Kumar et al., 2018). Amino acid sequences of CYP 3 clan in *D. melanogaster* were selected using Flybase (Öztürk-Çolak et al., 2024). Entire amino acid sequences of the enzymes in the housefly (*Musca domestica*), yellow fever mosquito (*Aedes aegypti*), silkmoth (*Bombyx mori*), western honey bee (*Apis mellifera*), red flour beetle (*Tribolium castaneum*), pea aphid (*Acyrthosiphon pisum*), and Nevada termite (*Zootermopsis nevadensis*) were obtained from the National Center for Biotechnology Information database (Sayers et al., 2022) using full length amino acid sequences of *D. melanogaster* proteins as queries. The protein names and GenBank accession numbers are listed in Table S7.

## 3. Results

### 3.1. Identification of CYPs highly expressed in the CA during early metamorphosis

We first investigated CYP enzymes that are highly expressed in the CA. In *D. melanogaster*, the CA is located within the RG, a composite endocrine organ (Zhang et al., 2022). We therefore analyzed previously published RNA sequencing data of the RGs from larvae at the wandering stage (Roger et al., 2023), when JH titers are known to be high (Sliter et al., 1987). We identified 12 CYP genes highly expressed in the RG (Table S4), including four ecdysteroid biosynthetic enzymes that are highly expressed in the prothoracic gland (Niwa and Niwa, 2026), which also constitutes the RG.

We next compared the expression levels of these CYP genes in the RG using qRT-PCR between control and CAX larvae. As the *Gal4* drivers previously used to induce CAX are also expressed in the salivary glands (Liu et al., 2009; Riddiford et al., 2010), we used a more CA-specific driver, *jhamt-LexA* (Ohhara et al., 2018; Fig. S1A), to ablate the CA by expressing pro-apoptotic genes (Ren et al., 2018; Fig. S1B). Among these genes, *Cyp6g2* and *Cyp6a13* showed high expression in the RG of control larvae as compared to CAX larvae (Fig. 2A), indicating that these genes are highly expressed in the CA, similar to *jhamt*. Absolute quantification of the gene expression revealed that *Cyp6g2* is expressed at a level more than 20 times higher than that of *Cyp6a13* in the control RG (Fig. S2). *In situ* hybridization further confirmed the expression of these genes in the CA, although the signal of the *Cyp6a13* antisense probe was weaker than that of *jhamt* or *Cyp6g2* due to its lower expression in the CA (Fig. 2B).

**Figure 2.**
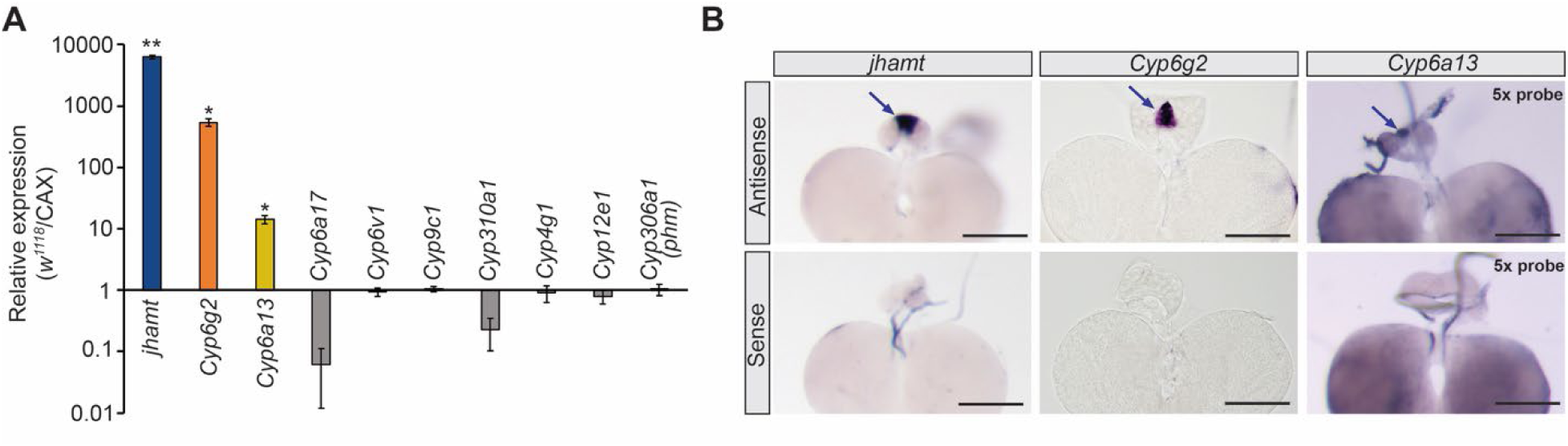
Cytochrome P450 genes highly expressed in the larval CA. (**A**) Relative gene expression levels in the ring gland (RG) of control (*w^1118^*) as compared to corpus allatum (CA)-ablated (CAX) larvae. *jhamt* and cytochrome P450 genes highly expressed in the RG of wandering larvae were quantified. High relative expression levels indicate CA-specific expression. Expression levels were normalized to a reference gene, *rp49*, in the same cDNA samples and shown on a log scale. Values are the means ± standard error (*n* = 3). \**p* < 0.05, \*\**p* < 0.01 (Student’s *t*-test). (**B**) *In situ* hybridization of *jhamt*, *Cyp6g2*, and *Cyp6a13* in the larval RG. Due to the lower expression of *Cyp6a13*, five times more RNA probe was used. The CA is indicated by arrows. Scale bars: 100 µm.

### 3.2. Combined mutations of the enzymes cause metamorphic defects similar to JH receptor mutants

We next generated null mutant alleles (Fig. S3) and analyzed the mutant flies to investigate contributions of these enzymes to *Drosophila* development. We also used our newly generated *Met*, *gce* double mutant line (*Met^45B^*, *gce^2C1^*) as a JH signaling-deficient control throughout this study, in order to exclude the potential effects of the intact coding sequence of the *Met^27^* allele (Wilson and Ashok, 1998) used in the previously reported *Met*, *gce* double mutant (Abdou et al., 2011).

Consistent with previous studies on JH signaling-deficient flies (Liu et al., 2009; Riddiford et al., 2010; Abdou et al., 2011), our new CAX flies died at different stages of metamorphosis (Fig. 3A). Based on this, we observed and classified developmental lethality of the mutant flies into the following five categories: larval lethal, prepupal lethal (PPL; characterized by incomplete air bubble translocation and/or head eversion), pupal lethal (PL), early adult lethal (EAL; defined as death within 24 hours after eclosion without wing expansion), and normal development. While almost all control flies (*w^1118^*) developed normally (Fig. 3B), most *Met*, *gce* double mutant flies died during pupation, with some showing the pupal lethality. Many CAX flies reached the pupal stage but eventually died during adult development or shortly after eclosion. *jhamt* single mutant flies showed mild phenotypes, as previously reported (Wen et al., 2015; Jia et al., 2024); most of these flies became adults, although about half of them died shortly after eclosion. Single mutants of *Cyp6g2* or *Cyp6a13* did not exhibit significant developmental defects, whereas combining them with the *jhamt* mutation showed much more severe phenotype, similar to *Met*, *gce* double mutants. Most *jhamt*, *Cyp6g2* double mutant flies died during or after pupation. *jhamt*, *Cyp6a13* double mutants showed slightly milder phenotypes than *jhamt*, *Cyp6g2* double mutants, although they were still significantly more severe than *jhamt* single mutant flies. *Cyp6g2*, *Cyp6a13* double mutants also showed mild lethal phenotypes. *jhamt*, *Cyp6g2*, *Cyp6a13* triple mutants showed severe phenotypes similar to *jhamt*, *Cyp6g2* double mutants. Altogether, these findings suggest that Jhamt, Cyp6g2, and Cyp6a13 function as JH synthetic enzymes and may have overlapping functions, as single mutants exhibited only mild phenotypes. Moreover, tissue specific knockdown using a *Gal4* driver expressed in the CA also suggested the JH synthetic activity of these enzymes in the CA (Fig. 3C). Single knockdowns of the CYPs did not cause significant developmental lethality. However, combined knockdown of *jhamt* and *Cyp6g2* or *jhamt* and *Cyp6a13* induced severe developmental defects, similar to the corresponding mutant combinations. Furthermore, the severity of developmental phenotypes of CAX and the mutant flies correlated with expression levels of a JH response gene, *Kr-h1* (Fig. 3D). Flies with mild developmental phenotypes (CAX; *jhamt*^-^; and *Cyp6g2*^-^, *Cyp6a13*^-^) exhibited moderate reduction of *Kr-h1* expression compared to *w^1118^* control, while those with more severe phenotypes (*Met^45B^*, *gce^2C1^*; *jhamt*^-^, *Cyp6g2*^-^; *jhamt*^-^, *Cyp6a13*^-^; and *jhamt*^-^, *Cyp6g2*^-^, *Cyp6a13*^-^) showed further reduction of *Kr-h1* expression levels. This strong correlation was confirmed by the correlation analysis (Fig. 3E), further suggesting the involvement of these enzymes in JH biosynthesis.

**Figure 3.**
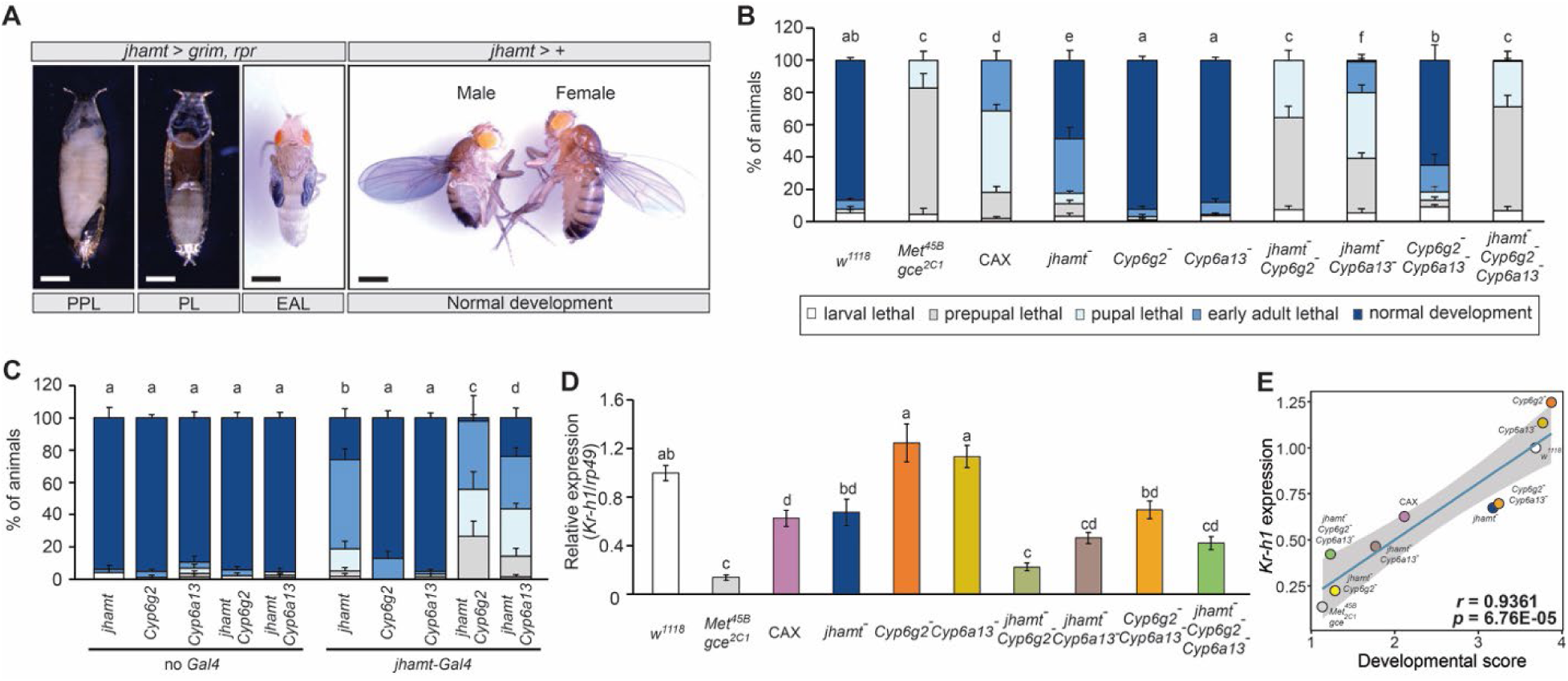
Lethal phenotypes of CA-ablated flies and JH-related mutants. (**A**) Representative images of the lethal phenotypes of corpus allatum (CA)-ablated (CAX) flies induced by *jhamt-LexA*-driven expression of *grim* and *reaper* (*rpr*). CAX flies died at different stages of metamorphosis. (**B**) Developmental lethality observed in CAX flies and homozygous mutants of JH-related genes. Receptor double mutants and combined mutants of *jhamt* with *Cyp6g2* or *Cyp6a13* exhibited severe lethality. Values are the means ± standard error. *n* = 3 (*Met^45B^*, *gce^2C1^*) or *n* = 6 (other mutants). Different letters above the bars indicate statistically significant differences (*p* < 0.05) between groups (Chi-square test with Bonferroni correction). (**C**) CA-specific knockdown of enzymes induced by *jhamt-Gal4*. Combined knockdown of *jhamt* with *Cyp6g2* or *Cyp6a13* exhibited severe lethality. Values are the means ± standard error (*n* = 3–4). Different letters above the bars indicate statistically significant differences (*p* < 0.05, Chi-square test with Bonferroni correction). (**D**) Relative expression levels of *Kruppel homolog 1* (*Kr-h1*) in CAX and mutant flies at 0 h after puparium formation. Expression levels were normalized to a reference gene, *rp49*, in the same cDNA samples. Flies with severe lethality tend to show lower expression levels. Values are the means ± standard error (*n* = 4–7). Different letters above the bars indicate statistically significant differences (*p* < 0.05, Tukey’s honestly significant difference test). (**E**) Correlation between developmental lethality and *Kr-h1* expression levels at 0 hour after puparium formation. Developmental scores were calculated based on lethal stages. A strong correlation (*r* = 0.9361) was observed between lethality and *Kr-h1* expression levels across various genotypes.

We also analyzed the pupal body size of CAX and the mutant flies (Fig. S4A). Previous studies have shown that JH-deficient flies exhibit a smaller body size along with slight developmental delay (Riddiford et al., 2010; Mirth et al., 2014). Consistent with this, *Met*, *gce* double mutant, CAX, *jhamt*, *Cyp6g2* double mutant, and *jhamt*, *Cyp6g2*, *Cyp6a13* triple mutant flies were significantly smaller than *w^1118^* control flies in both sexes. Indeed, correlation analyses confirmed a strong correlation between developmental lethality and pupal body size reduction (Fig. S4B and C). In contrast, pupariation timing was not significantly altered, although mutants with high lethality tended to exhibit slightly delayed pupariation timing (Fig. S4D).

### 3.3. Sesquiterpenoid feeding rescues developmental defects of the enzyme mutants

In order to confirm that developmental defects of CAX flies and the mutants were caused by impaired JH biosynthesis, we applied JHs and their precursors to the third instar larvae (Fig. 4A-H). Control flies showed no developmental defects when treated with the sesquiterpenoids (Fig. 4A). As expected, no rescue was observed in *Met*, *gce* double mutants (Fig. 4B), as they lack the molecular machinery to respond to JH. In contrast, developmental defects of CAX flies were rescued by all terpenoids tested, particularly MF, JH III, and JHB3 (Fig. 4C). These control experiments demonstrate that exogenous sesquiterpenoid application can successfully compensate for endogenous JH functions without any harmful side effects during development. It should be noted that MF, like JH III and JHB3, can also activate JH receptors, although its activity is weaker than JH III and JHB3 (Jindra et al., 2015). The enzyme mutants were also rescued by MF, JH III, and JHB3 in a dose-dependent manner (Fig. 4D-H), suggesting that the lethality observed in the mutant flies was indeed due to reduced JH production. Conversely, application of the JH precursor, FA, hardly rescued the enzyme mutants, suggesting that these enzymes function downstream of FA in the JH biosynthetic pathway (Fig. 1). Interestingly, significant rescue of the developmental lethality in CAX flies was observed when the higher concentration (5 ppm) of FA was applied, potentially suggesting the existence of an alternative JH biosynthetic machinery outside the CA.

**Figure 4.**
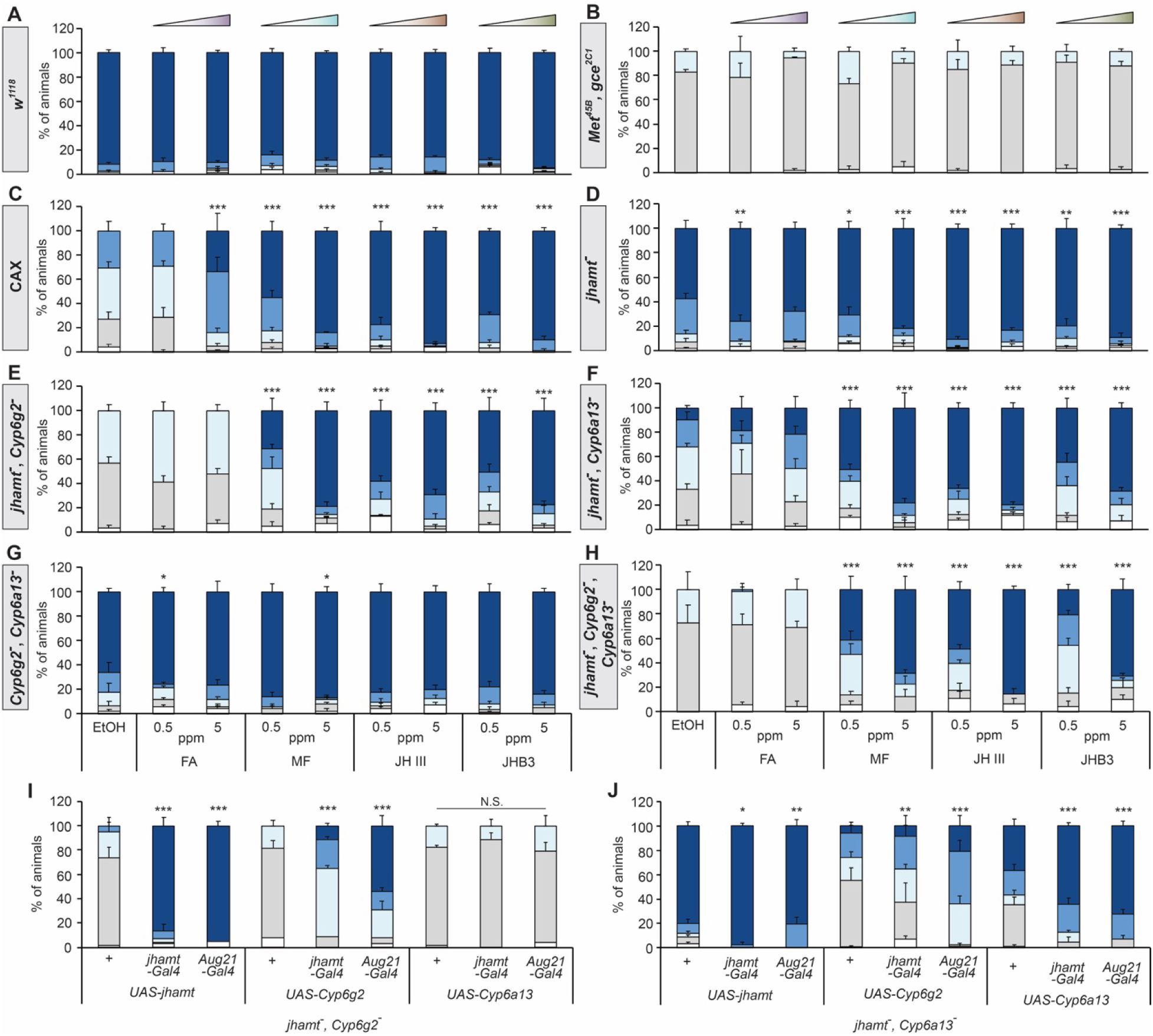
Rescue of CA-ablated flies and JH-related mutants. (**A-H**) Rescue of developmental defects observed in control (*w^1118^*), corpus allatum (CA)-ablated (CAX) flies, and homozygous mutants of JH-related genes by application of farnesoic acid (FA), methyl farnesoate (MF), juvenile hormone III (JH III), or juvenile hormone III bisepoxide (JHB3). Larvae were transferred onto the diet containing 0.5 or 5 ppm of one of the sesquiterpenoids at 48 hours after hatching and raised until eclosion. MF, JH III, and JHB3 rescued lethality of most genotypes, whereas FA had little effect. Values are the means ± standard error (*n* = 3–6). \**p* < 0.05, \*\**p* < 0.01, \*\*\**p* < 0.001 (Chi-square test with Bonferroni correction *vs* application of the solvent (EtOH) alone). (**I, J**) Rescue of *jhamt*, *Cyp6g2* (I) or *jhamt*, *Cyp6a13* (J) double mutants by CA-specific overexpression of *jhamt*, *Cyp6g2*, or *Cyp6a13*. *Cyp6g2* overexpression partially rescued *jhamt*, *Cyp6a13* double mutants, but *Cyp6a13* overexpression did not rescue *jhamt*, *Cyp6g2* double mutants. Values are the means ± standard error (*n* = 3–6). \*\**p* < 0.01, \*\*\**p* < 0.001 (Chi-square test with Bonferroni correction *vs* control flies without a *Gal4* driver). N.S., not significant (*p* > 0.05).

We also performed rescue experiments of *jhamt*, *Cyp6g2* and *jhamt*, *Cyp6a13* double mutants by overexpressing each enzyme in the CA (Fig. 4I and J). Consistent with their functions in the CA, overexpression of the enzymes in their corresponding mutants significantly rescued the developmental defects. Importantly, *Cyp6g2* overexpression significantly rescued *jhamt*, *Cyp6a13* double mutants, whereas *Cyp6a13* overexpression did not rescue *jhamt*, *Cyp6g2* double mutants. These results suggest that Cyp6g2 can compensate for Cyp6a13 in JH biosynthesis in the CA, whereas Cyp6a13 plays a more limited role.

### 3.4. Contribution of the enzymes to reproduction

During metamorphosis, the CA and corpora cardiaca (CC) complex migrates toward the ventral ganglion and eventually settles on the esophagus in *D. melanogaster* (Dai and Gilbert 1991; Fig. S1A). Although *Cyp6a13* expression levels were significantly lower than the other enzymes (∼1.5% compared to *jhamt* expression), transcripts of all the three enzyme-encoding genes were detectable in the adult CA/CC complex (Fig. 5A-C). Among the three enzyme-encoding genes, *jhamt* expression increased within 24 hours after eclosion, potentially reflecting the elevated JH III titer after eclosion (Reiff et al., 2015; Sugime et al., 2017).

**Figure 5.**
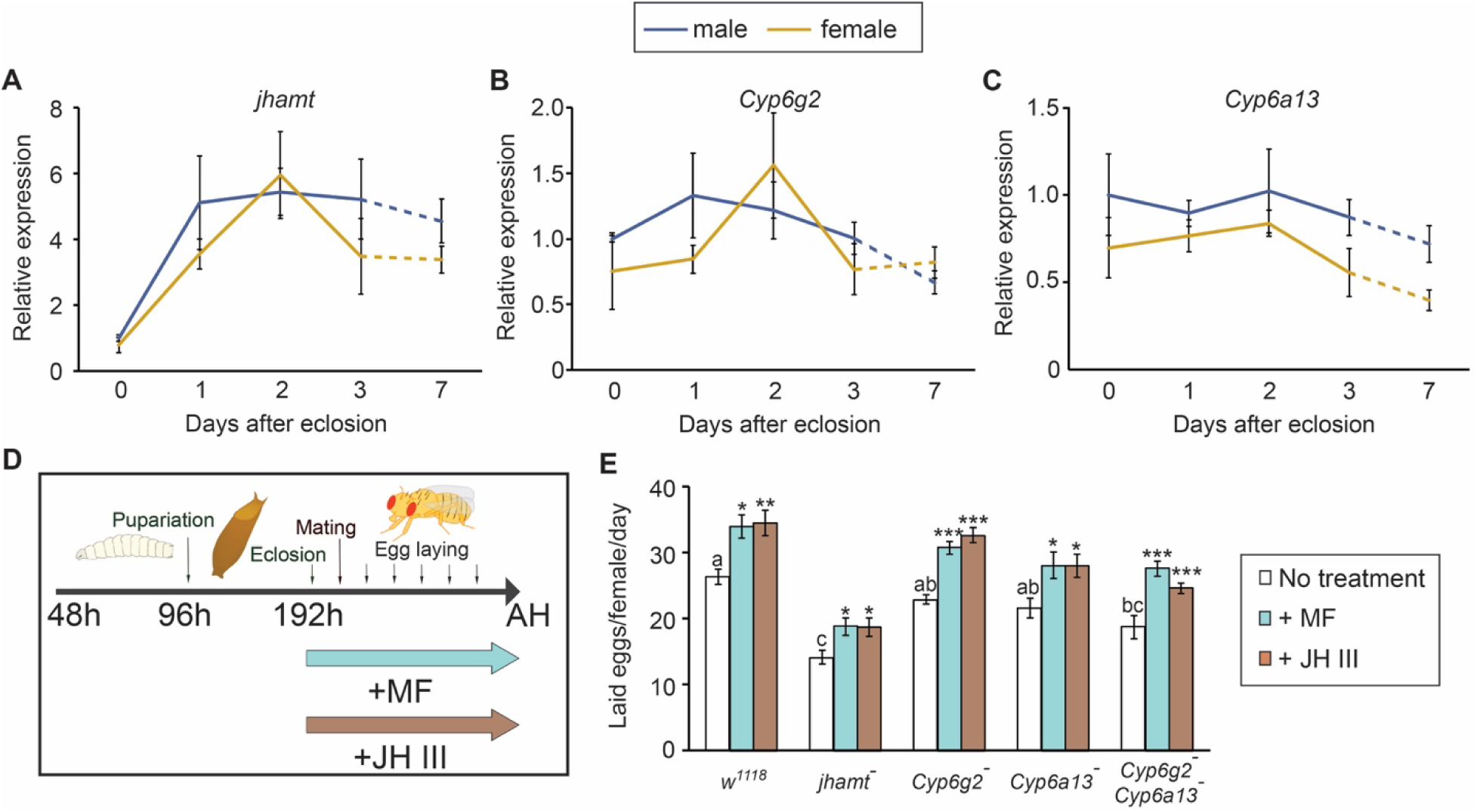
Contributions of the CA-expressed enzymes to reproduction. (**A–C**) Changes in gene expression of *jhamt* (A), *Cyp6g2* (B), and *Cyp6a13* (C) in the corpus allatum/corpora cardiaca complex of the adults expressing *jhamt-LexA*-driven *LexAop-mCD8::GFP*. Expression levels were normalized to levels at day 0 after eclosion (within 6 hours). Values are the means ± standard error (*n* = 3–4). (**D**) Schematic diagram of the fecundity assay. Mutants were reared normally until eclosion. Eclosed adults were then maintained with or without methyl farnesoate (MF) or juvenile hormone III (JH III). Mutant females were mated with *w^1118^* males 24 hours after eclosion. The number of eggs laid was counted every 24 hours from 3 to 7 days after eclosion. *Drosophila* illustrations were obtained from TogoTV (© 2016 DBCLS TogoTV, CC-BY-4.0 https://creativecommons.org/licenses/by/4.0/deed.ja). (**E**) Number of eggs laid by the mutant females. The numbers of eggs laid by untreated females were compared across genotypes using Tukey’s honestly significant difference test. Different letters above the bars indicate statistically significant differences among different genotypes (*p* < 0.05). The numbers of eggs laid by MF- or JH III-treated females were compared to that of untreated females of the same genotype using Dunnett’s test (\**p* < 0.05, \*\**p* < 0.01, \*\*\**p* < 0.001). *jhamt* mutant and *Cyp6g2*, *Cyp6a13* double mutant females laid significantly fewer eggs than *w^1118^* control. Sesquiterpenoid treatment enhanced egg production in all genotypes tested. Values are the means ± standard error (*n* = 6–12).

Given the functions of JHs as gonadotropic factors in adult insects including *D. melanogaster* (Marchal et al., 2014; Roy et al., 2018; Santos et al., 2019; Smykal et al., 2014; Song et al., 2014), we next measured the number of eggs laid by the mutant flies to assess the contributions of these enzymes to reproduction (Fig. 5D). *jhamt* single mutant and *Cyp6g2*, *Cyp6a13* double mutant females laid significantly fewer eggs compared to *w^1118^* control (Fig. 5E). On the other hand, neither *Cyp6g2* nor *Cyp6a13* single mutants displayed statistically significant reduction of the number of laid eggs. These results suggest that Cyp6g2 and Cyp6a13 may compensate for each other in JH production in adult females. Treatment with MF or JH III after eclosion significantly increased the fecundity of females of all genotypes, further suggesting the importance of JH signaling in reproduction in female flies.

### 3.5. Cyp6g2 and Cyp6a13 are responsible for epoxidation of JH precursors

To further investigate enzymatic activities *in vitro*, we modified the sesquiterpenoid extraction described in Sarro et al. (2021) to enable simultaneous extraction of MF, JH III, and JHB3 for GC-MS and LC-MS analyses. The modified hexane extraction method recovered nearly 100% of FA, MF, JH III, and JHB3 (Fig. S5). Using this method, we conducted an enzymatic conversion assay in HEK293 cells (Fig. 6A). Overexpression of *jhamt* induced methylation of FA as expected from previous studies (Niwa et al., 2008), while its co-expression with either *Cyp6g2* or *Cyp6a13* did not enhance MF production (Fig. 6B). In contrast, co-expression of *Cyp6g2* with *jhamt* enhanced conversion of FA into JH III beyond the basal level of JH III production observed in *jhamt* single overexpression (Fig. 6C), consistent with the reported epoxidase activity of Cyp6g2 (Jia et al., 2024). Importantly, co-expression of *Cyp6a13* with *jhamt* also promoted conversion of FA into JH III (Fig. 6C), suggesting that it encodes an epoxidase similar to Cyp6g2.

**Figure 6.**
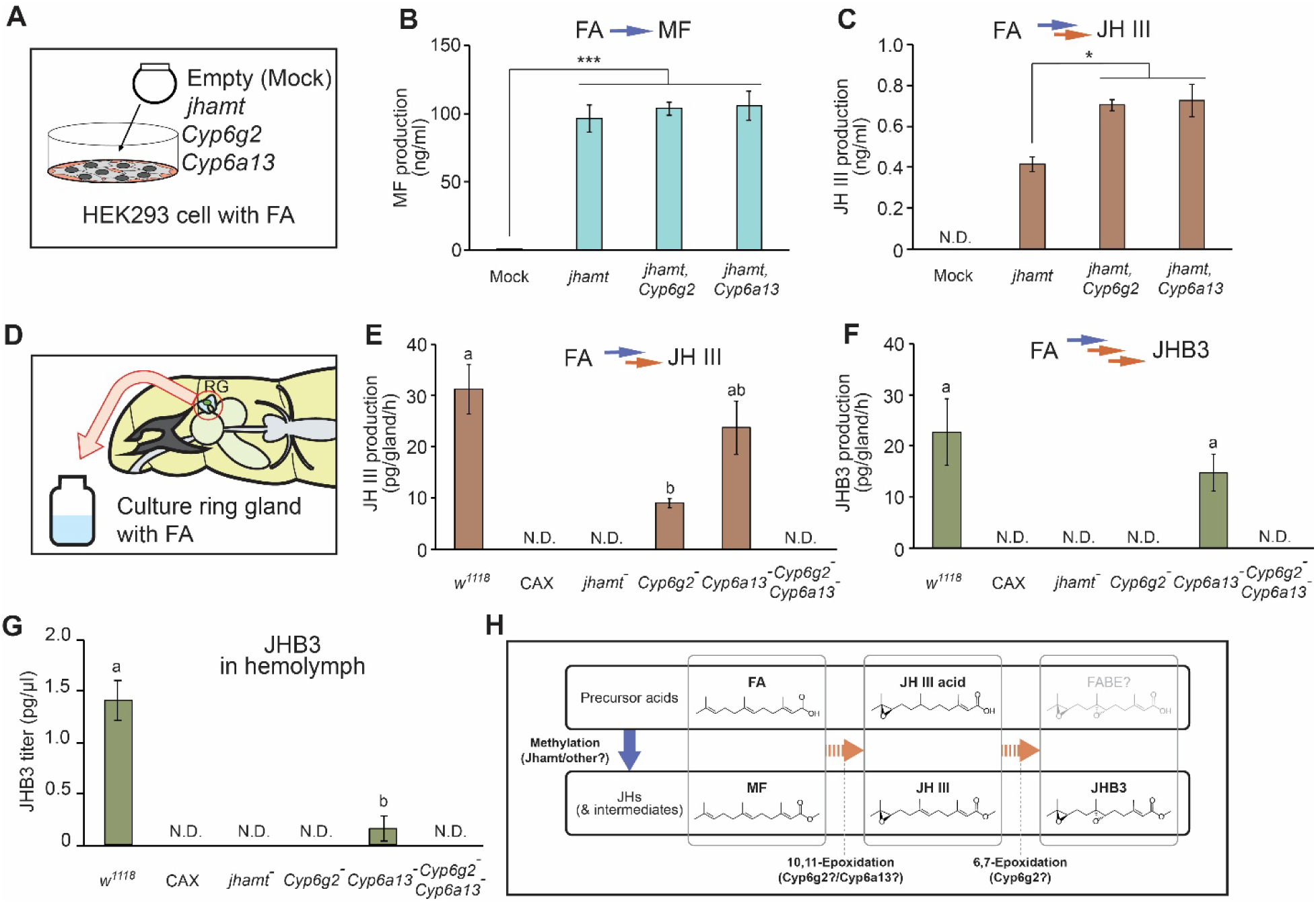
Functional analyses of the CA-expressed enzymes. (**A**) Schematic diagram of the enzymatic conversion assay in HEK293 cells. (**B, C**) Conversion of farnesoic acid (FA) into methyl farnesoate (MF) (B) and juvenile hormone III (JH III) (C) in HEK293 cells ectopically expressing the CA-expressed enzymes. Jhamt catalyzes methylation, while Cyp6g2 and Cyp6a13 catalyze epoxidation to synthesize JH III. Values are the means ± standard error (*n* = 3–4). \**p* < 0.05, \*\*\**p* < 0.001 (Tukey’s honestly significant difference test). N.D., not detected. (**D**) Schematic diagram of *ex vivo* culture of the ring glands from wandering larvae. (**E, F**) Conversion of FA into JH III (E) and juvenile hormone III bisepoxide (JHB3) (F) in cultured ring glands. JH production in the *jhamt* mutant ring glands was drastically reduced. The *Cyp6g2* mutant ring glands produced only a small amount of JH III. While JH production in the *Cyp6a13* mutant ring glands was not significantly reduced, it was abolished in the *Cyp6g2*, *Cyp6a13* double mutant ring glands. Different letters above the bars indicate significant differences (*p* < 0.05, Tukey’s honestly significant difference test). Values are the means ± standard error (*n* = 4–12). N.D., not detected. (**G**) JHB3 titers in the hemolymph of wandering larvae of control (*w^1118^*), corpus allatum (CA)-ablated (CAX) flies, and the enzyme mutants. The hemolymph JHB3 titer was significantly reduced in *Cyp6a13* mutant larvae, and it was undetectable in CAX, *jhamt* mutant, *Cyp6g2* mutant, and *Cyp6g2*, *Cyp6a13* double mutant larvae. MF and JH III were below detection limits. Different letters above the bars indicate significant differences (*p* < 0.05, Tukey’s honestly significant difference test). Values are the means ± standard error (*n* = 4–7). N.D., not detected. (**H**) Proposed model of the JH biosynthetic pathway in the *Drosophila* CA.

Contrary to our expectations, JHB3 was not detected under any condition in the enzymatic conversion assay in HEK293 cells. We speculate that an unknown cofactor present in the CA may be necessary for JHB3 biosynthesis *in vivo*. We therefore conducted *ex vivo* culture of the larval RGs to assess enzymatic activity of CYPs in the CA (Fig. 6D). JH III and JHB3 were detected in the RGs from control larvae cultured with FA (Fig. 6E and F), confirming the methylation and epoxidation activities in the cultured RGs. In the *Cyp6g2* mutant RG, JH III production was significantly decreased (Fig. 6E), and JHB3 production was below the detection limit (Fig. 6F). These results indicate that Cyp6g2 acts as an epoxidase that can produce both JH III and JHB3. Although the RGs from the *Cyp6a13* single mutant larvae did not show statistically significant reduction of JH III or JHB3 production, JH III was not produced in the cultured RGs from the *Cyp6g2*, *Cyp6a13* double mutant larvae (Fig. 6E). This further suggests that Cyp6a13 epoxidizes JH precursors to synthesize JH III.

Lastly, hemolymph JH levels were analyzed to confirm the JH biosynthetic activities of the enzymes *in vivo*. In the hemolymph of wandering larvae, we could only detect JHB3, as the reported amounts of larval JH III are far below our detection limit (∼5 pg/µL hemolymph; Jones et al., 2013; Wen et al., 2015; Barton et al., 2024). Compared to control, the hemolymph level of JHB3 was significantly reduced in the *Cyp6a13* single mutant larvae, and it was not detected in CAX larvae and the other mutants tested (Fig. 6G). Overall, our results indicate that both Cyp6g2 and Cyp6a13 function as epoxidases in the CA to produce JHs *in vivo*.

## 4. Discussion

In the present study, we showed that two CYP enzymes, Cyp6g2 and Cyp6a13, likely function as epoxidases for JH biosynthesis in the *Drosophila* CA. Higher dipteran insects, known as *Cyclorrhapha*, have lost the canonical sesquiterpenoid epoxidase CYP15 in their genomes (Dermauw et al., 2020; Smykal and Dolezel, 2023). Instead, they employ CYP6 enzymes as epoxidases: for example, Cyp6a1 has been identified as an MF epoxidase in the housefly *Musca domestica* (Andersen et al., 1997). A recent study showed that Cyp6g2 is a major epoxidase for JH production in *D. melanogaster* (Jia et al., 2024). The present study now adds *Drosophila* Cyp6a13 to the list of CYP6 epoxidases. Considering that the RGs isolated from wandering *Cyp6g2*, *Cyp6a13* double mutant larvae did not produce detectable levels of JHs (Fig. 6E and F), it is likely that Cyp6g2 and Cyp6a13 are the only epoxidases that contribute to JH production in the *Drosophila* CA, at least during the wandering stage.

The relatively mild developmental defects seen in *Cyp6g2*, *Cyp6a13* double mutants are likely due to the weak but significant JH receptor-activating capacity of MF (Jindra et al., 2015; Wen et al., 2015). Consistent with this, previous reports suggest that MF is circulating in the *Drosophila* hemolymph during development (Jones et al., 2013; Wen et al., 2015). Further studies on this JH-like function of MF are required to evaluate the significance of epoxidation in JH biosynthesis in *Drosophila*.

JHB3 is a unique JH species in higher dipteran insects that have lost CYP15 epoxidases (Smykal and Dolezel, 2023). It is therefore likely that JHB3 is a specific product of CYP6 epoxidases. Although we failed to detect JHB3 production by the CYP6 enzymes in HEK293 cells, this result must be interpreted with caution, as human cells may lack specific components (such as cofactors) required for full CYP6 activities. Consistent with this, although we detected JH III biosynthetic activities of the CYP6 enzymes in this assay, less than 1% of MF produced by JHAMT was converted to JH III (Fig. 6B and C), further highlighting the limitation of this heterologous expression system.

Given that the *Cyp6g2* mutant RGs do not synthesize detectable amounts of JHB3 *ex vivo*, we infer that Cyp6g2 is the only epoxidase that can synthesize JHB3 in *D. melanogaster*. Although the hemolymph level of JHB3 was drastically reduced in *Cyp6a13* single mutant larvae, this may reflect reduced production of JH III, the precursor of JHB3. Indeed, the small amount of JH III produced by the *Cyp6g2* mutant RGs disappears in *Cyp6g2*, *Cyp6a13* double mutant RG cultures, suggesting that Cyp6a13 is at least involved in JH III production in the RG. The inferred functions of the CYP6 enzymes are summarized in Fig. 6H.

In *D. melanogaster*, it is known that the hemolymph JHB3 titer is much higher than that of JH III (Wen et al., 2015; Ramirez et al., 2020). This is consistent with our hemolymph JH analysis, in which we only detected JHB3. It is therefore interesting that comparable amounts of JH III and JHB3 were produced in the RGs cultured *ex vivo*. A potential explanation for this is that JH-degrading enzymes such as JH esterases and JH epoxide hydrolases (Kamita and Hammock, 2010; Siegel et al., 2025) may preferentially degrade JH III over JHB3. Although a previous *in vitro* study showed that *Drosophila* JH esterase has similar affinity for both JH III and JHB3 (Campbell et al., 1998), another study suggested that high concentrations of JH esterase preferentially degrades JH III when bound to its binding proteins (Touhara et al., 1995). Since JHB3 does not bind to lipophorin, a major JH-binding protein (Trowell et al., 1994), JH esterase may be less effective at degrading JHB3 as compared to JH III. Elucidating molecular mechanisms behind the high titer of JHB3 in the *Drosophila* hemolymph may help us better understand the physiological significance of this higher dipteran-specific JH species.

*D. melanogaster* Cyp6g2 and Cyp6a13, as well as *M. domestica* Cyp6a1, are distributed within the insect CYP3 clan. Although *Drosophila* Cyp6a13 and *Musca* Cyp6a1 are located within the same cluster, CYP3 enzymes of these two *Cyclorrhapha* species are greatly expanded within this subclade (Fig. S6). Taken together, these observations suggest that their JH biosynthetic functions evolved independently. Enzymes in the CYP3 clan (such as Cyp6 and Cyp9 enzymes) are highly divergent in insects, and many are known to have xenobiotic-detoxifying activities (Nauen et al., 2022). *Drosophila* Cyp6g2 and *Musca* Cyp6a1 are also known for their detoxification roles (Feyereisen, 1999; Giang et al., 2025). Many other CYP3 clan enzymes catalyze epoxidation of their target compounds as well (Feyereisen, 1999). It is therefore possible that other enzymes in this clan are also capable of epoxidizing sesquiterpenoids, but only those expressed in the CA have obtained JH biosynthetic functions. We could not find any other CYP enzymes specifically expressed in the CA, suggesting that other *Drosophila* enzymes do not contribute to JH biosynthesis *in vivo*, at least during the wandering stage. In contrast, it remains possible that other CYPs function as JH synthases in peripheral tissues, as some studies suggest that JH biosynthesis occurs in other organs, such as the adult gut (Rahman et al., 2017; Shianiou et al., 2023). These studies showed that *jhamt* expression in the gut is responsible for maintaining cellular growth. Notably, *Cyp6a13* is expressed not only in the CA, but also in peripheral tissues, according to Flybase expression data (Öztürk-Çolak et al., 2024). It is therefore possible that Cyp6a13 acts with Jhamt for JH biosynthesis in peripheral tissues.

In summary, our current work revealed that two CYP6 enzymes, Cyp6g2 and Cyp6a13, function cooperatively to synthesize JH in the *Drosophila* CA. Further investigations of their differential functions, along with identification of additional JH biosynthetic enzymes, are expected to help us better understand both species-specific and conserved features of JH signaling in insects.

## Conflict of interest

The authors declare no competing interest.

## Acknowledgement

We thank Vienna *Drosophila* Resource Center, Bloomington *Drosophila* Stock Center (supported by NIH P40 OD018537), National Institute of Genetics, T. Lee, M. E. Adams, and M. B. O’Connor for fly stocks; FlyBase for providing curated *Drosophila* genome information; Metabolomics core at the University of California, Riverside for JHB3 determination; *Drosophila* Genomics Resource Center (supported by NIH 2P40 OD010949) and National Institute of Genetics for vectors and cDNA clones. This study was supported by the Japan Society for the Promotion of Science (JSPS) KAKENHI 21H02521 to Y.O., a Postdoctoral Fellowship for Research Abroad from JSPS to N.O., the Naito Foundation Subsidy for Dispatch of Young Researchers Abroad to N.O., an NIH grant R01 AI171032 from NIAID to N.Y., and an NIH grant R35 GM153331 from NIGMS to N.Y.

## Author contributions

DF: conceptualization, methodology, investigation, formal analysis, writing – original draft, writing – review and editing. YO: methodology, investigation, funding acquisition, writing – review and editing. NO: investigation, funding acquisition, writing – review and editing. HC: generation of *Met*, *gce* mutant flies, writing – review and editing. KM: methodology for sesquiterpenoid determination, writing – review and editing. NY: conceptualization, funding acquisition, project administration, supervision, writing – review and editing.

## Notes

### Competing Interest Statement

The authors have declared no competing interest.

### Summary of Updates

Additional discussion was added to clarify the significance and limitations of the study; additional references were incorporated.

