## Supplementary Information for "Two cytochrome P450 epoxidases mediate juvenile hormone biosynthesis in *Drosophila melanogaster*"

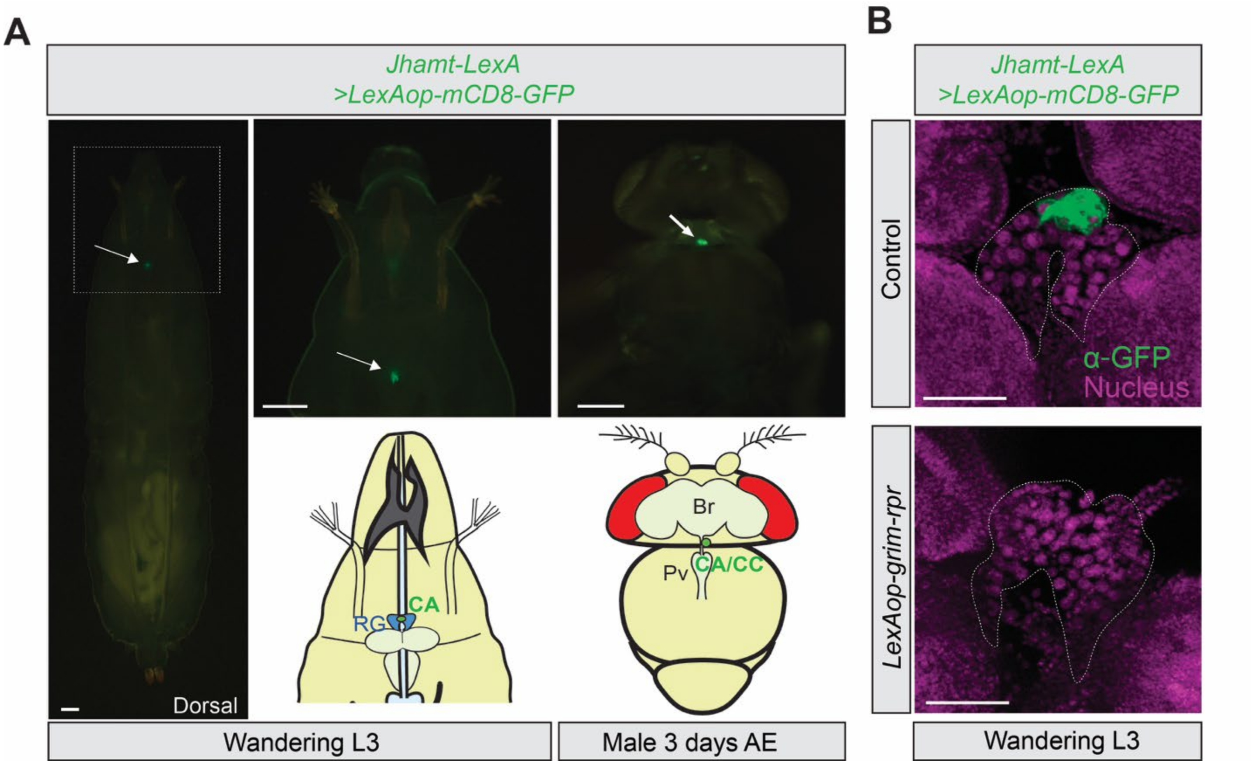


**Figure S1. Visualization of *Jhamt-LexA* expression patterns.**

(**A**) Expression patterns of *Jhamt-LexA* visualized by *LexAop-mCD8::GFP* expression. The corpus allatum (CA) is indicated by arrows. *Jhamt-LexA* is specifically expressed in the CA of wandering third instar (L3) larvae and adults. In the image of a whole larva, the anterior region indicated by the square is enlarged. Scale bars, 200 µm. RG, ring gland. AE, after eclosion. Br, brain. Pv, proventriculus. CA/CC, corpus allatum/corpora cardiaca complex. (**B**) Apoptosis induction in the CA by CA-specific overexpression of *grim* and *reaper* (*rpr*). CA cells labeled with GFP were lost in CAX larvae. The ring gland is outlined with white dotted lines. GFP expression in the CA was detected by immunostaining. Scale bars, 50 µm.

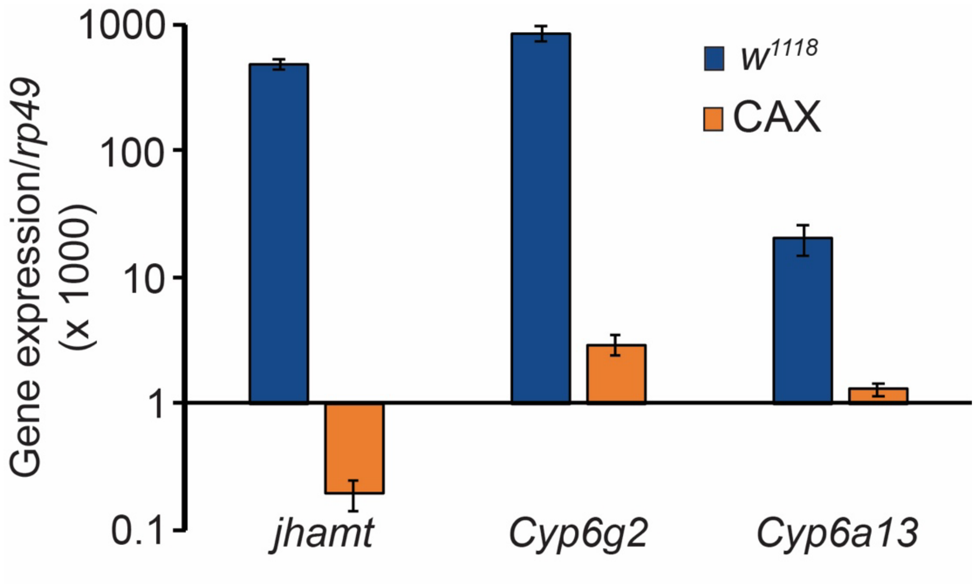


**Figure S2. Absolute quantification of gene expression in the ring gland.**

Absolute quantification of *jhamt*, *Cyp6g2*, and *Cyp6a13* expression in the ring gland of control ($w^{1118}$) and corpus allatum-ablated (CAX) larvae. Expression levels were normalized to a reference gene, *rp49*, in the same cDNA samples and shown on a log scale. A standard curve for each gene was generated using serial dilutions of a plasmid containing the target sequence. All target genes showed proportional amplification relative to the amount of the template plasmid (*jhamt*: efficiency = 95.7%, $R^2$ = 0.999; *Cyp6g2*: efficiency = 90.1%, $R^2$ = 0.995; *Cyp6a13*: efficiency = 119.5%, $R^2$ = 0.995; *rp49*: efficiency = 93.9%, $R^2$ = 0.999). Values are the means ± standard error ($n$ = 3).

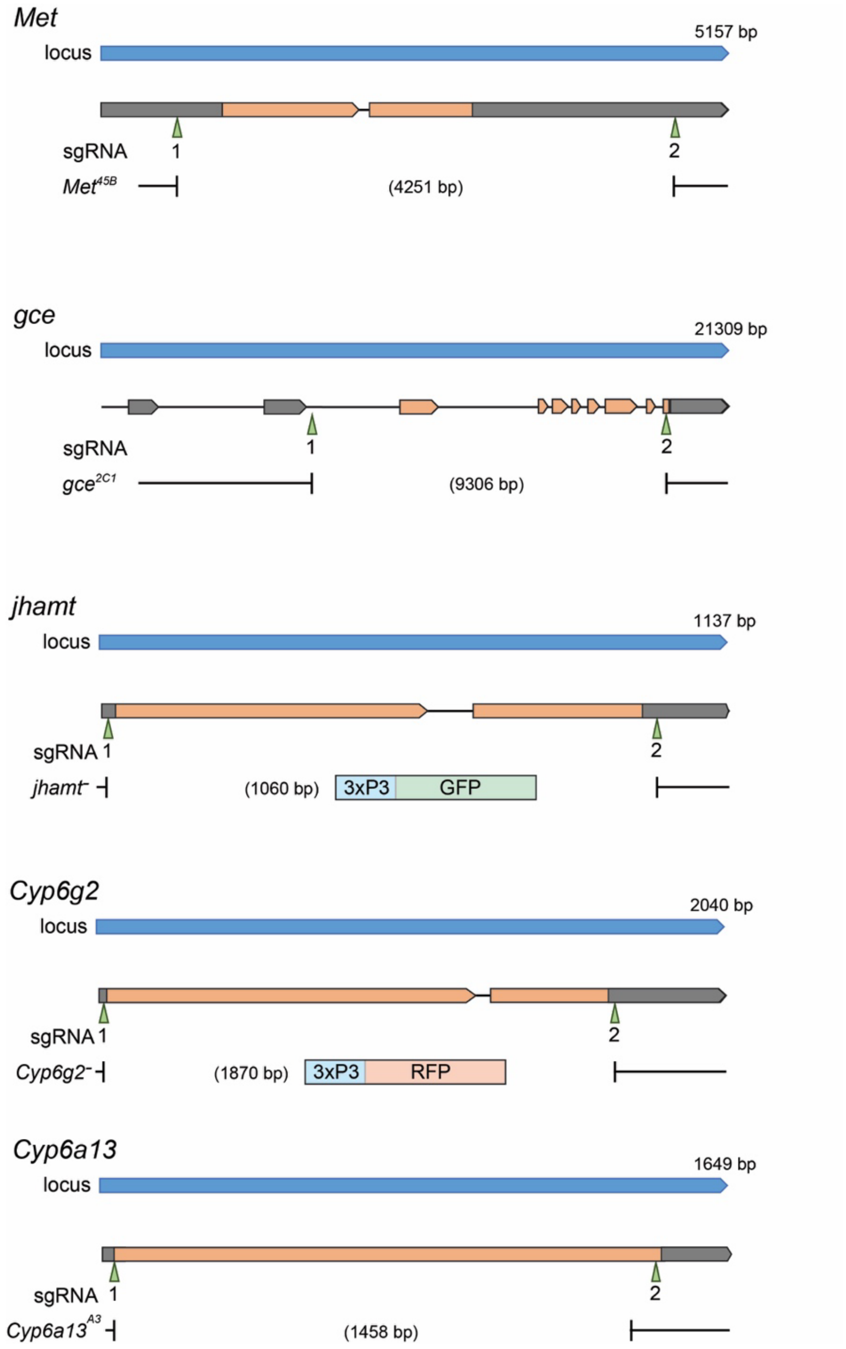


**Figure S3. Mutagenesis of JH-related genes.**

CRISPR-Cas9-mediated mutagenesis was performed using two single guide RNAs (sgRNAs) per target gene to delete most or all of the coding sequences (indicated in orange). A 3xP3-GFP sequence was inserted into the *jhamt* target site, and a 3xP3-RFP sequence was inserted into the *Cyp6g2* target site via homologous recombination.

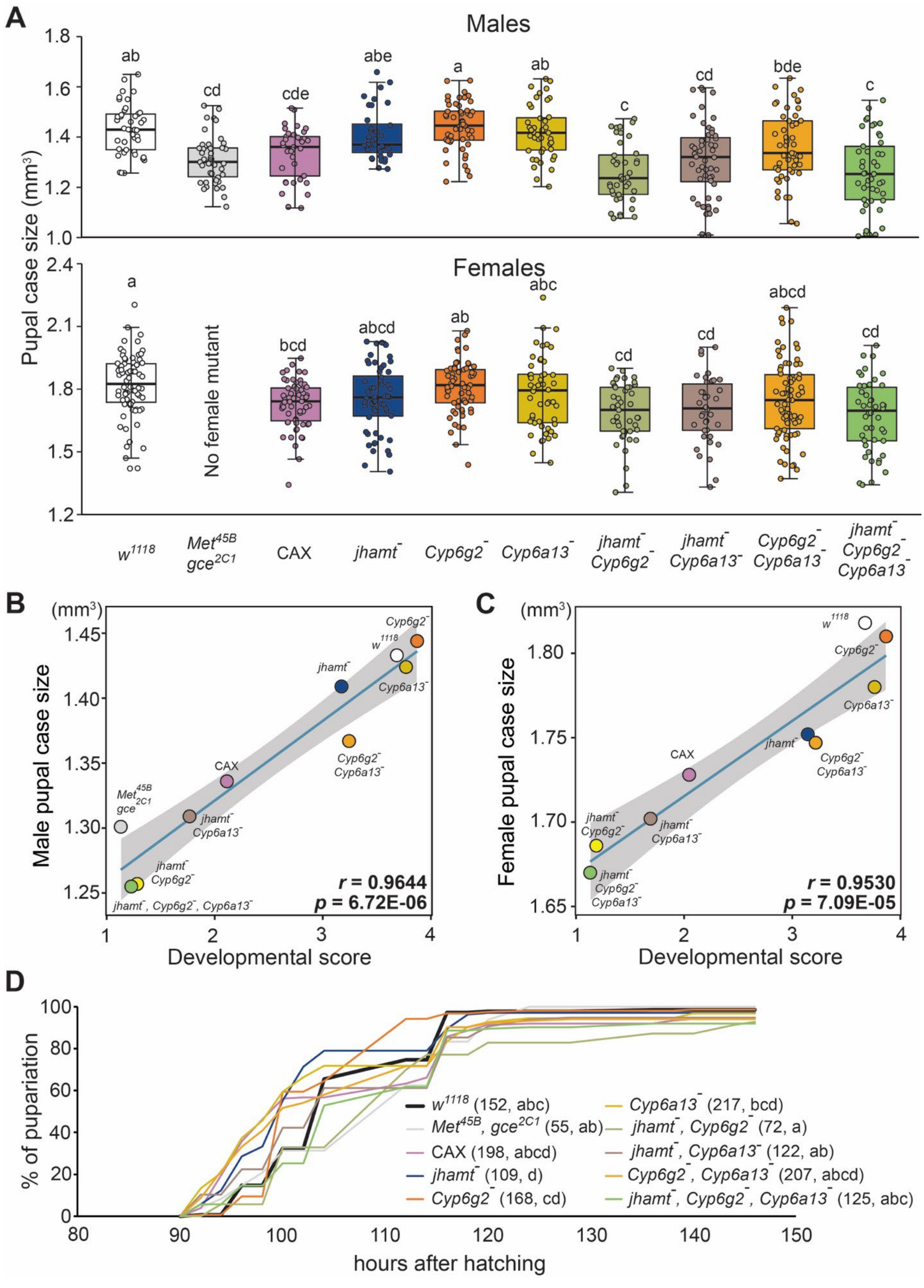

A
Males
Females
Pupal case size (mm³)
No female mutant
w1118
Met45B gce2C1
CAX
jhamt⁻
Cyp6g2⁻
Cyp6a13⁻
jhamt⁻ Cyp6g2⁻
jhamt⁻ Cyp6a13⁻
Cyp6g2⁻ Cyp6a13⁻
jhamt⁻ Cyp6g2⁻ Cyp6a13⁻
B
Male pupal case size
Developmental score
r = 0.9644
p = 6.72E-06
C
Female pupal case size
Developmental score
r = 0.9530
p = 7.09E-05
D
% of pupariation
hours after hatching
w1118 (152, abc)
Met45B, gce2C1 (55, ab)
CAX (198, abcd)
jhamt⁻ (109, d)
Cyp6g2⁻ (168, cd)
Cyp6a13⁻ (217, bcd)
jhamt⁻, Cyp6g2⁻ (72, a)
jhamt⁻, Cyp6a13⁻ (122, ab)
Cyp6g2⁻, Cyp6a13⁻ (207, abcd)
jhamt⁻, Cyp6g2⁻, Cyp6a13⁻ (125, abc)

**Figure S4. Developmental phenotypes of CA-ablated flies and JH-related mutants.**

(**A**) Pupal case size of CAX and the homozygous mutant males and females. Pupal case sizes of each genotype and sex are shown as scatter and box plots. Animals with severe lethal phenotypes tend to show smaller body sizes. *Met*/*gce* double mutant females were not available as these genes are on the X chromosome. $n$ = 36–73. Different letters indicate statistically significant differences ($p < 0.05$) between groups (Tukey's honestly significant difference test). (**B, C**) Correlation between the developmental score and male (B) or female (C) body size. Developmental scores were calculated based on lethal stages. Strong positive correlations ($r >$ 0.95, Pearson's correlation coefficient) were observed in both cases. (**D**) Pupariation timing after hatching was analyzed in CAX larvae and JH-related mutants. Most larvae entered the prepupal stage between 90 and 120 hours after hatching. Numbers in parentheses indicate the numbers of larvae analyzed. Different letters in parentheses indicate statistically significant differences between genotypes ($p < 0.05$, log-rank test).

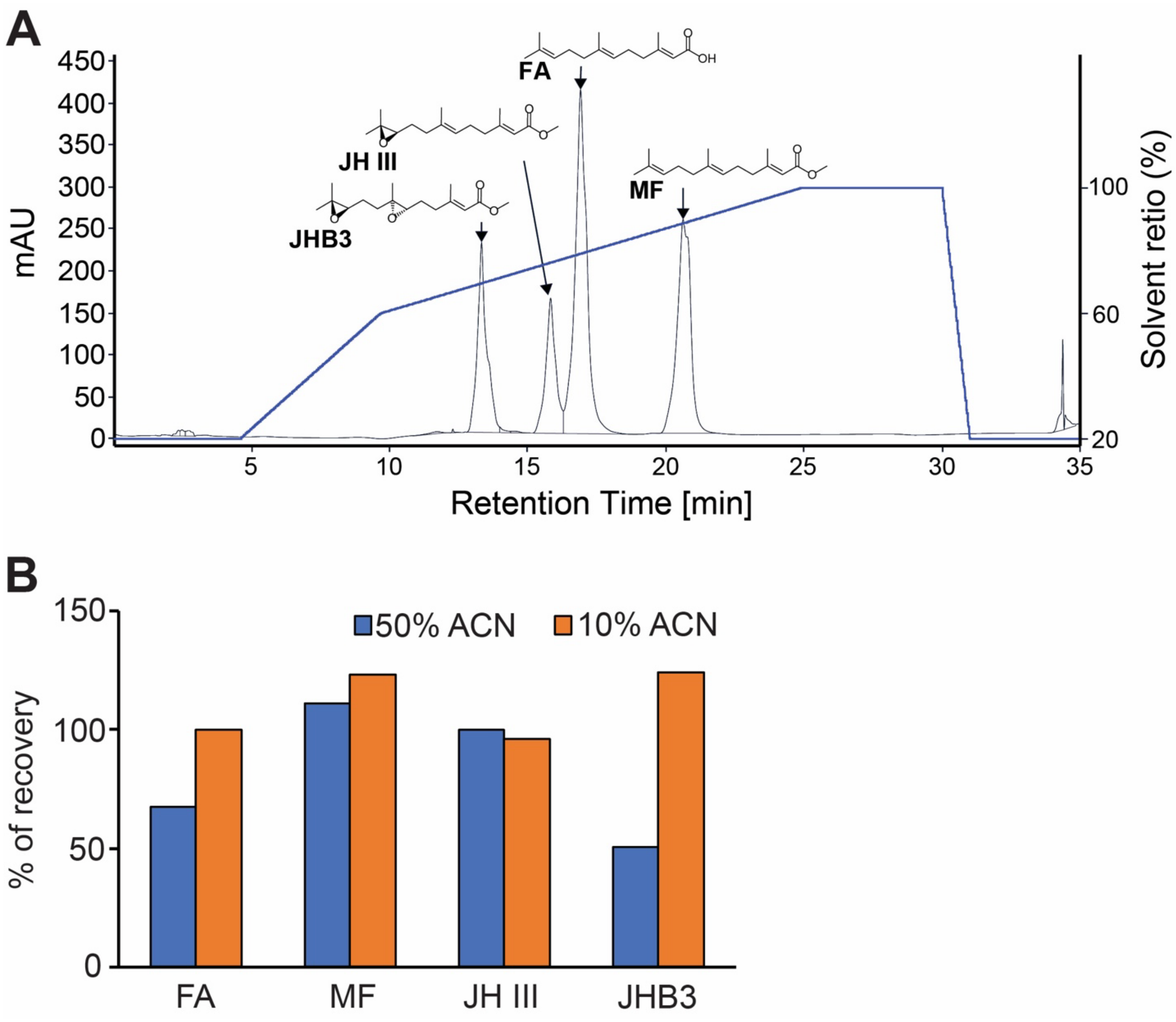


**Figure S5. Simultaneous extraction of sesquiterpenoids.**

(**A**) Chromatogram of the sesquiterpenoid mixture containing 1 µg each of authentic standards. (**B**) Recovery rate of each compound following hexane extraction from solutions containing different concentrations of acetonitrile (ACN). Note that recovery rates exceeding 100% are due to partial evaporation of methanol during concentration with the CentriVap concentrator. FA, farnesoic acid; MF, methyl farnesoate; JH III, juvenile hormone III; JHB3, juvenile hormone III bisepoxide.

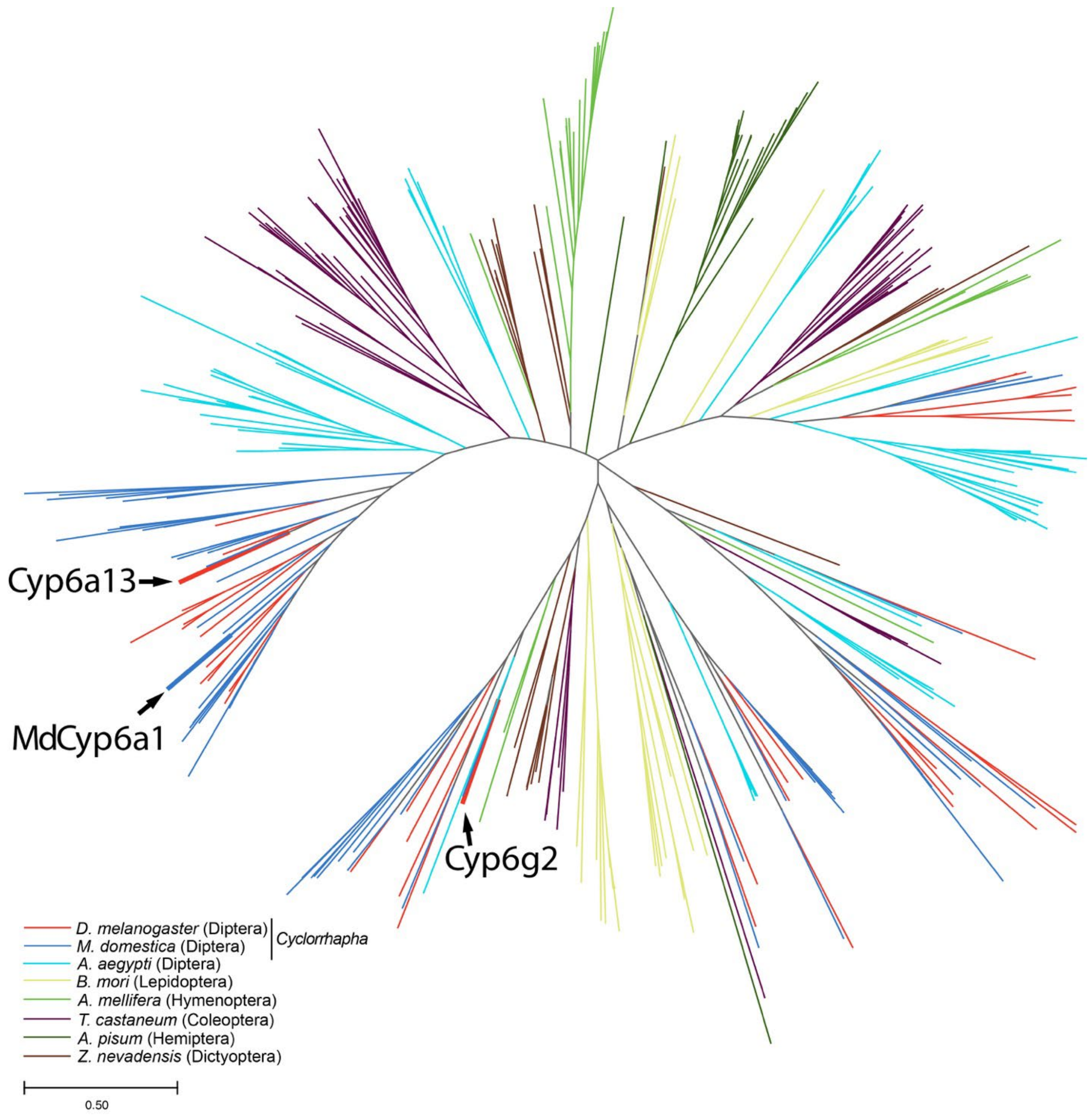


**Figure S6. Phylogenetic analysis of enzymes in the insect CYP3 clan.**

Unrooted maximum-likelihood phylogenetic tree of CYP3 clan enzymes in *Drosophila melanogaster*, *Musca domestica, Aedes aegypti, Bombyx mori*, *Apis mellifera*, *Tribolium castaneum*, *Acyrthosiphon pisum*, and *Zootermopsis nevadensis*. The scale bar indicates an evolutionary distance of 0.5 amino acid substitutions per site. Accession numbers of the enzymes analyzed are listed in Table S7.

**Table S1. *Drosophila* strains used in this study.**

| Organisms: Strains | Source | Identifier |
|---|---|---|
| *D. melanogaster*: $w^{1118}$ | Bloomington *Drosophila* Stock Center | BDSC: 5905 |
| *D. melanogaster*: *jhamt-LexA* | Ohhara et al., 2018 | |
| *D. melanogaster*: *LexAop-mCD8::GFP* | Bloomington *Drosophila* Stock Center | BDSC: 56173 |
| *D. melanogaster*: *LexAop-grim, rpr* | Obtained from Tzumin Lee (University of Michigan) | N/A |
| *D. melanogaster*: *nos-Cas9* | National Institute of Genetics | CAS-0002 |
| *D. melanogaster*: *U6-Cyp6a13 gRNA* | This paper | N/A |
| *D. melanogaster*: *jhamt-Gal4* | Obtained from Michael E. Adams (University of California, Riverside) | N/A |
| *D. melanogaster*: *Aug21-Gal4* | Obtained from Michael B. O'Connor (University of Minnesota) | N/A |
| *D. melanogaster*: $Met^{45B}$ | This paper | N/A |
| *D. melanogaster*: $gce^{2C1}$ | This paper | N/A |
| *D. melanogaster*: $jhamt^{-}$ | This paper | N/A |
| *D. melanogaster*: $Cyp6g2^{-}$ | This paper | N/A |
| *D. melanogaster*: $Cyp6a13^{-}$ | This paper | N/A |
| *D. melanogaster*: *UAS-jhamt RNAi* | Vienna *Drosophila* Resource Center | VDRC: 103958 |
| *D. melanogaster*: *UAS-Cyp6g2 RNAi* | Vienna *Drosophila* Resource Center | VDRC: 8131 |
| *D. melanogaster*: *UAS-Cyp6a13 RNAi* | Vienna *Drosophila* Resource Center | VDRC: 4019 |
| *D. melanogaster: UAS-jhamt* | This paper | N/A |
| *D. melanogaster: UAS-Cyp6g2* | This paper | N/A |
| *D. melanogaster: UAS-Cyp6a13* | This paper | N/A |

**Table S2. Plasmid constructs used in this study.**

| Recombinant DNA | Source | Identifier |
|---|---|---|
| cDNA AT13581 (*jhamt*) | *Drosophila* Genomics Resource Center | DGRC: 11840<br>FlyBase: FBcl0041099 |
| cDNA IP03155 (*Cyp6g2*) | *Drosophila* Genomics Resource Center | DGRC: 1379871<br>FlyBase: FBcl0343443 |
| cDNA LD25139 (*Cyp6a13*) | *Drosophila* Genomics Resource Center | DGRC: 2249<br>FlyBase: FBcl0175511 |
| *pcDNA-jhamt* | This paper | N/A |
| *pcDNA-Cyp6g2* | This paper | N/A |
| *pcDNA-Cyp6a13* | This paper | N/A |
| *pBFv-U6.2* | National Institute of Genetics | N/A |
| *pBFv-U6.2B* | National Institute of Genetics | N/A |
| *pUAST-jhamt* | This paper | N/A |
| *pUAST-Cyp6g2* | This paper | N/A |
| *pUAST-Cyp6a13* | This paper | N/A |

**Table S3. Primers and oligonucleotides used in this study.**

| **Primers for qRT-PCR** | | | |
|---|---|---|---|
| Gene Name | CG Number | Forward (5'-3') | Reverse (5'-3') |
| *jhamt* | CG17330 | TTTCTTGAGCGAATGCCTGC | AGGAGTCTTGCGAGCATAGGC |
| *Cyp6g2* | CG8859 | AGGATACGGACGTACAGCAG | GCATGAACTCCAGTGACTCG |
| *Cyp6a13* | CG2397 | ATGGAATTGGGTGAGGAGGG | AGAAGGACATGGTGGTGGAG |
| *Cyp6a17* | CG10241 | ATTTCCCAAGTTATCGCGCC | CGCTTTTCCTTCGTCCGTAG |
| *Cyp6v1* | CG1829 | TATCCCTCCACGATTCAGCC | TCAGGATCGTCACTATCGCC |
| *Cyp9c1* | CG3616 | TGGGGCGGACTCAAAGTAAT | TCGATACTTCATGGCACCGA |
| *Cyp310a1* | CG10391 | TATTGAACGTGTTTGCCGCA | TCGGTGCATAGCCAGTAGAG |
| *Cyp4g1* | CG3972 | CGTCCAGACATCTACCCCAA | GTCCAGCGCTAAAGGGAATG |
| *Cyp12e1* | CG14680 | CCACGTAACGCCAAGCTTTA | GTCCACGGCAAAGTCATCTG |
| *phm* | CG6578 | CTTCTGGTCCAATCCCGACT | ATTCGTTTCTCTTTCCGCGG |
| *Kr-h1* | CG45974 | GAATACGACATAACAGCC | CGATTTCCGTGAATATGTTCT |
| *rp49* | CG7939 | AGCTGTCGCACAAATGGCGCAAGC | TTGAATCCGGTGGGCAGCATGTGG |

| **Oligonucleotides for generating gRNAs** | | | |
|---|---|---|---|
| Allele | Target ID | Forward (5'-3') | Reverse (5'-3') |
| $Met^{45B}$ | 1 | CTTCGAAGGAGAGTGAGCGAGAGA | AAACTCTCTCGCTCACTCTCCTTC |
| | 2 | CTTCGTTGGTGATGGGCGCAGAGA | AAACTCTCTGCGCCCATCACCAAC |
| $gce^{2C1}$ | 1 | CTTCGACTGAATCTGTCGGTGCAA | AAACTTGCACCGACAGATTCAGTC |
| | 2 | CTTCGTGATGGAGAGCGCCCGATC | AAACGATCGGGCGCTCTCCATCAC |
| $Cyp6a13^{-}$ | 1 | CTTCGAGTACCAGGAGCGTCAGCA | AAACTGCTGACGCTCCTGGTACTC |
| | 2 | CTTCGGAGGGAGTTCATTTGAGGA | AAACTCCTCAAATGAACTCCCTCC |

| **Primers for screening CRISPR mutants** | | |
|---|---|---|
| Allele | Forward (5'-3') | Reverse (5'-3') |
| $Met^{45B}$ | GCCACCAGCAACAGCAACAACAAT | CGCTGGGGAATTAGCATTGGACTT |
| $gce^{2C1}$ | GTAGCAGCCCAAGGACTATACCAT | CCATCAGTTGACCGAACGAGAAGT |
| *Cyp6a13-* | TGTGTCAGCGCTTCAAGTTC | ACATATTTCGCGCTGTCAGG |

| **Primers for generating *in situ* hybridization probes** | | |
|---|---|---|
| Gene Name | | Sequence (5'-3') |
| *jhamt* | Forward | GTAATACGACTCACTATAGGGCACTCCTGGATGTGGGTTCAG |
| | Reverse | GTGTATTTAGGTGACACTATAGGCATAGGCCACCACCAATTT |
| *Cyp6g2* | Forward | GTAATACGACTCACTATAGGGCGTGCCAACAGTTTCACGGAT |
| | Reverse | GTGTATTTAGGTGACACTATAGCCACCTTCGCCTGGAGATAT |
| *Cyp6a13* | Forward | GTAATACGACTCACTATAGGGCACGGATATCAACCTGCGGAT |
| | Reverse | GTGTATTTAGGTGACACTATAGAGAAGGACATGGTGGTGGAG |

**Table S5. MS Settings to quantify each compound on the GC-MS.**

| Compound | Retention time (min) | Ion Polarity | Window (min) | Parent Mass | Product Mass | Collision Energy (V) |
|---|---|---|---|---|---|---|
| Citronellol | 5.87 | Positive | 0.2 | 81.0 | 79.1 | 10 |
| Methylfarnesoate | 10.12 | Positive | 0.2 | 114.1 | 83.1 | 10 |
| Juvenile hormone III | 11.01 | Positive | 0.2 | 85.1 | 59.1 | 10 |

**Table S6. MS Settings to quantify each compound on the LC-MS.**

| Compound | Retention time (min) | Ion Polarity | Window (min) | Parent Mass | Product Mass | Collision Energy (V) |
|---|---|---|---|---|---|---|
| Farnesoic acid | 14.23 | Negative | 1.0 | 235.2 | 98.2 | 12 |
| Juvenile hormone III | 13.44 | Positive | 1.1 | 267.2 | 235.0 | 12 |
| Juvenile hormone III-d3 | 13.41 | Positive | 0.9 | 270.2 | 153.2 | 20 |
| Juvenile hormone bisepoxide | 10.79 | Positive | 1.25 | 283.2 | 233.0 | 18 |
